# Macroevolutionary analysis of discrete traits with rate heterogeneity

**DOI:** 10.1101/2020.01.07.897777

**Authors:** Michael C. Grundler, Daniel L. Rabosky

## Abstract

Organismal traits show dramatic variation in phylogenetic patterns of origin and loss across the Tree of Life. Understanding the causes and consequences of this variation depends critically on accounting for heterogeneity in rates of trait evolution among lineages. Here, we describe a method for modeling among-lineage evolutionary rate heterogeneity in a trait with two discrete states. The method assumes that the present-day distribution of a binary trait is shaped by a mixture of stochastic processes in which the rate of evolution varies among lineages in a phylogeny. The number and location of rate changes, which we refer to as rate-shift events, are inferred automatically from the data. Simulations reveal that the method accurately reconstructs rates of trait evolution and ancestral character states even when simulated data violate model assumptions. We apply the method to an empirical dataset of mimetic coloration in snakes and find elevated rates of trait evolution in two clades of harmless snakes that are broadly sympatric with dangerously venomous New World coral snakes, recapitulating an earlier analysis of the same dataset. Although the method performed well on many simulated data sets, we caution that overall power for inferring heterogeneous dynamics of single binary traits is low.

## Introduction

Organismal traits evolve recurrently across the Tree of Life, and the frequency with which traits evolve varies widely among clades. Among extant amniotes, for example, viviparity has evolved at least 100 times but only a single origin of it (in mammals) occurs outside squamate reptiles (Blackburn 1982, 1985). Similarly, sugar-secreting glands known as extrafloral nectaries have arisen hundreds of times among plants, mainly within the legumes, but not once among gymnosperms (Weber and Keeler 2013). Identifying the causes and consequences of repeated convergence depends critically on accounting for heterogeneity in rates of trait evolution. Ancestral trait reconstructions reveal evidence of convergence and can identify other traits enabling the repeated evolution of a convergent trait (Maddison 1990; de Queiroz and Rodríguez-Robles 2006; Marazzi et al. 2012; Christin et al. 2013). However, inferences of ancestral states may be seriously misled by models that fail to account for rate heterogeneity (King and Lee 2015). Several evolutionary theories also link how quickly traits evolve to how quickly lineages diversify (Vermeij 1973; Stanley 1979), and the repeated evolution of a trait together with methods that model among-lineage variation in evolutionary rates allows for direct tests of this coupling (Rabosky et al. 2013; Igea et al. 2017).

Methods to account for evolutionary rate heterogeneity have grown in number and in sophistication over recent years (O’Meara 2006; Revell and Collar 2009; Eastman et al. 2011; Lloyd et al. 2012; Marazzi et al. 2012; Beaulieu et al. 2013; Landis et al. 2013; Rabosky et al. 2014; Uyeda and Harmon 2014), but much of this methodological progress has focused on continuous traits such as body mass or seed size. Phylogenetics deals extensively with rate heterogeneity in discrete character evolution because of the challenge it poses for inference of evolutionary relationships (e.g. Heath et al. 2012), and some of these methods carry over to the study of non-molecular traits. The “hidden rates” (Beaulieu et al. 2013) and “precursor” (Marazzi et al. 2012) models, first introduced to study variation in growth form and extrafloral nectary production in plants, are closely related to covarion models of nucleotide substitution (Fitch and Markowitz 1970; Galtier 2001; Penny et al. 2001). The “random local clock” model (Drummond and Suchard 2010) used to study variation in the rate of nucleotide substitution has also been used to study evolutionary rate variation in mimetic color pattern (Davis Rabosky et al. 2016) and reproductive mode parity (King and Lee 2015) in squamate reptiles.

In this paper, we describe a method for modeling evolutionary rate heterogeneity in a trait with two discrete states and implement it using the Bayesian Analysis of Macroevolutionary Mixtures (BAMM) framework (Rabosky 2014; Rabosky et al. 2014). The model discussed here is closely related to several existing phylogeny inference methods that model among-lineage substitution rate variation (Huelsenbeck 2000; Drummond and Suchard 2010) but differs in details of likelihood calculation and implementation. The general approach assumes that the present-day distribution of a binary state character is shaped by a mixture of stochastic processes in which the rate of evolutionary transition between the two states experiences shifts under a Poisson process across the branches of a phylogeny. The number and location of rate changes, which we refer to as rate-shift events, are inferred automatically from the data. Simulations reveal that the method accurately infers rates of evolution, the number and location of rate-shift events, and ancestral character states even when simulated data violate model assumptions.

## Materials & Methods

### Likelihood of a binary state character

We assume that each branch of a phylogeny evolves independently of the others and that the probability of a lineage transitioning to a character state different from its current state does not depend on its prior history of trait evolution (Pagel 1994). For a discrete character with two states, state 0 and state 1, trait evolution is modeled by a “forward” transition rate (denoted *q*_01_), which governs how frequently a lineage in state 0 changes to state 1, and a “reverse” transition rate (denoted *q*_10_), which governs how frequently a lineage in state 1 changes to state 0. For example, over a sufficiently small interval time Δ*t*, the probability of observing a transition from state 0 to state 1 is approximately *q*_01_Δ*t*.

To write down a likelihood function for estimating the transition rate parameters, we follow the approach of Maddison et al. (2007) and define *D*_*N*0_(*t*) to be the probability that lineage *N* evolves the distribution of character states observed among its descendants given that it is in state 0 at time *t*. We define *D*_*N*1_(*t*) analagously. Next, consider what can happen in the short interval of time between *t* and *t+h*, where *t+h* is closer to the root and where *h* is taken to be an interval of time small enough that the probability of more than one character state transition in the interval is negligible. There are only two possibilities. Either the lineage remains in the state that it was in at time *t+h* or it switches to the other state. We can therefore write *D*_*N*0_(*t* + *h*) and *D*_*N*1_(*t* + *h*) as functions of *D*_*N*0_(*t*), *D*_*N*1_(*t*), and the transition rate parameters *q*_01_ and *q*_10_,

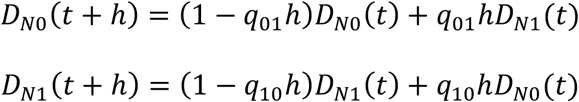

By rearranging and letting *h* approach zero we form two coupled differential equations that describe how these probabilities change through time,

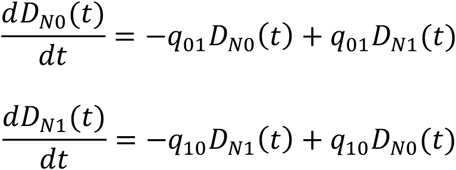

These can be solved (see appendix) to give the closed form solutions,

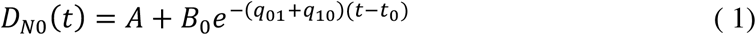

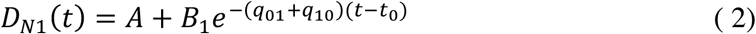

Where *t*_0_ is the initial time and,

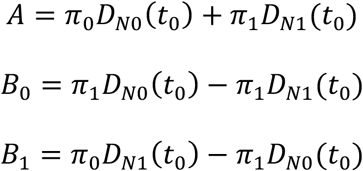

Where *π_i_* is the equilibrium frequency of state *i*. Equations (1) and (2) are evaluated for each branch of the phylogeny proceeding from the tips to the root in a post-order traversal. The quantities *D*_*N*0_(*t*_0_) and *D*_*N*1_(*t*_0_) are the initial conditions used to begin evaluation for each branch. If we have these values, we can compute the conditional likelihood of the data for a branch’s stem group by simply setting *t* equal to the time at the base (rootward) of the branch and *t*_0_ equal to the time at the head (tipward) of the branch and evaluating equations (1) and (2). At each internal node, we form a new set of initial conditions by multiplying the *D*_·0_ and *D*_·1_ at the base of the node’s left descendant branch with those of its right descendant. When we reach the root, *R*, of the tree *D*_*R*0_(*t_R_*) and *D*_*R*0_(*t_R_*) yield the probability of the data given the transition rate parameters and conditional on the root being in state 0 or state 1, respectively. To get the unconditional likelihood we must combine these two values, but doing so requires knowing the probability of the root being in state 0 or in state 1. We assume that the root is in state 0 or state 1 with probabilities implied by their conditional likelihoods and compute the full likelihood as 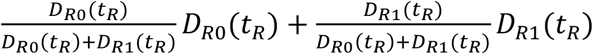. This is the same weighting scheme used in some models of trait-dependent speciation and extinction (FitzJohn et al. 2009). To begin computation, the initial conditions for each tip in the tree are set to *D*_·0_(0) = 1 and *D*_·1_(0) = 0 if the tip is in state 0 and vice versa if the tip is in state 1.

### Rate-shift model for a binary state character

At broad phylogenetic scales, it may be unreasonable to assume that the transition rates remain constant over all regions of phylogeny. We therefore allow regions of a phylogeny to belong to different macroevolutionary “rate-regimes”, which are independent sets of transition rate parameters that describe the evolution of a binary state character over the regions of phylogeny to which they pertain. Transitions between rate-regimes, which we refer to as “rate-shift events”, are assumed to occur along the branches of a phylogeny according to a compound Poisson process (Huelsenbeck 2000; Blanquart and Lartillot 2006; Rabosky 2014). To compute the likelihood of the data under a rate-shift model requires minimal modification of the process described in the previous section. Rather than traversing directly from a node to its ancestor when evaluating equations (1) and (2), we pause at each rate-shift event that occurs on the branch and use the values of (1) and (2) at that point as starting values for an additional evaluation of (1) and (2) under the new set of transition rates.

This model of rate variation is unsuitable when forward and reverse transition rates are asymmetric (Fig. 1). By proposing a new rate-shift event for every event of character state change and simply making one rate arbitrarily large and the other rate arbitrarily small it is possible to fit any data set with probability 1 (when conditioned on the occurrence of the rate-shift events). This is because we can guarantee with probability 1 the origin and persistence of a derived character state by making the transition rate toward the derived character state arbitrarily large and the reverse rate arbitrarily small. In such a scenario, rate-shift events become decoupled from broad-scale among-lineage variation in the rate of trait evolution. Our implementation of an asymmetric version of this rate-shift model yielded results consistent with these expectations for several empirical datasets. For this reason, the default implementation in BAMM constrains forward and reverse transition rates to be identical. All analyses presented below use this symmetric implementation.

**Figure 1.**
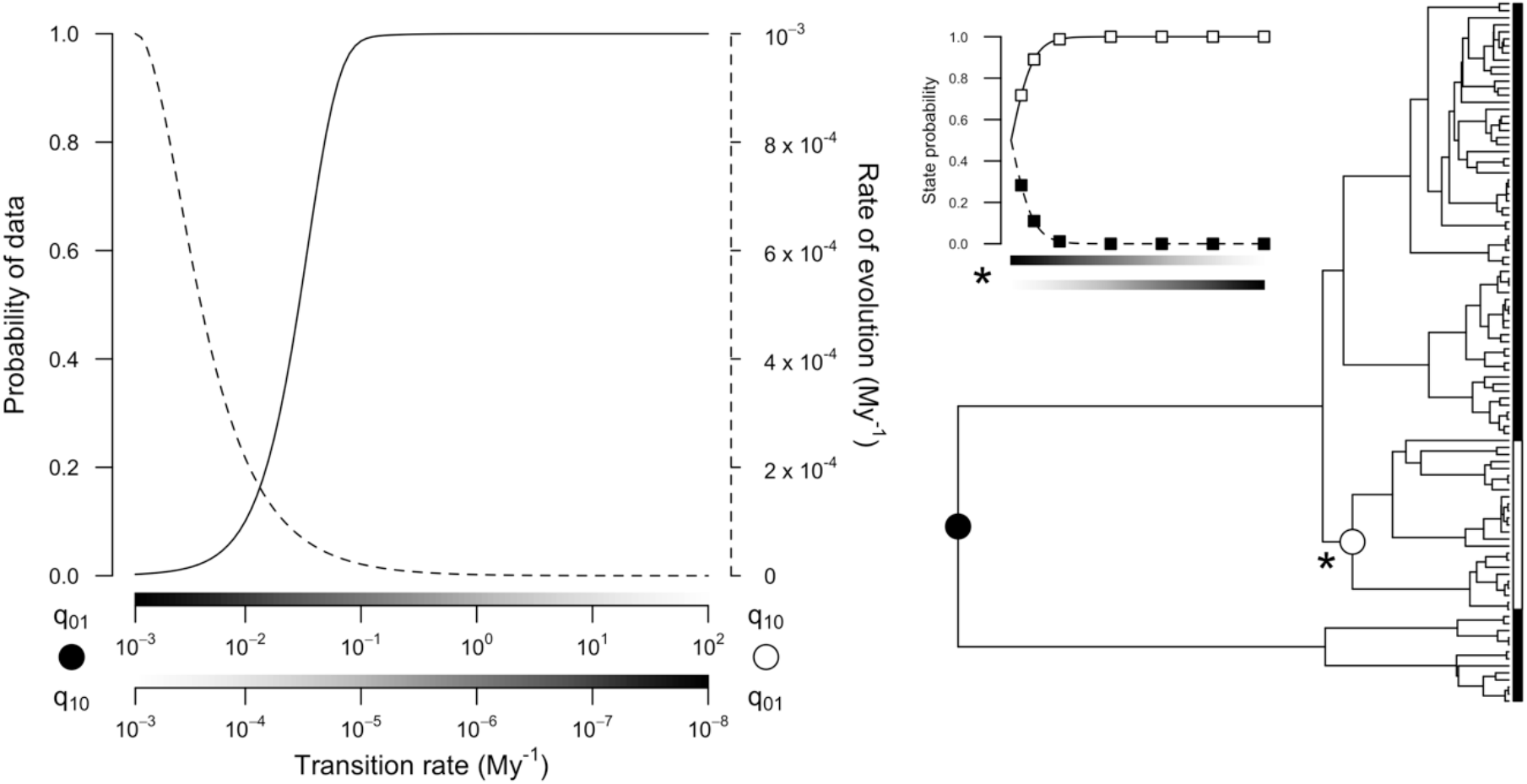
A worked example showing why asymmetric Markov models are unsuitable for BAMM-type rate-shift models. A single rate-shift event (denoted by a white circle) is placed along the branch subtending the clade fixed for character state 0. The *x*-axes in the left plot depict transition rates for this rate-shift and for the root event (denoted by a black circle). Note that they run in opposite directions, e.g. the upper *x*-axis is *q*_01_ for the root event but is *q*_10_ for the rate-shift event. As the asymmetry in transition rates is increased the probability of the data (denoted by the solid line) rises to 1 while the overall rate of character evolution (denoted by the dashed line) falls to 0. Simultaneously, as the rate of character evolution falls to 0 the asymmetry in transition rates causes the prior probability of being in one or the other character state to rise to 1, e.g. the inset plot denoted by an asterisk depicts the probability of being in state 0 or state 1 as a function of the asymmetric transition rates of the rate-shift event. Thus, at the extreme a “rate-shift” event simply introduces a second way to observe a stochastic event of character state change and does not correspond to among-lineage heterogeneity in rates of character evolution.

### Implementation

The rate-shift model described above is implemented in the Bayesian software program BAMM using reversible-jump Markov Chain Monte Carlo simulation (Rabosky 2014). Briefly, BAMM assumes that the number of rate-shift events on a phylogeny is drawn from a Poisson distribution with a rate parameter that is itself drawn from an exponential distribution. This formulation implies that the number of rate-shifts is drawn from a geometric distribution and that the expected number of rate-shifts, denoted by Λ, is simply the mean of the exponential hyperprior placed on the Poisson prior (Mitchell and Rabosky 2017). We extend the BAMM implementation for binary data by placing an exponential prior on the transition rate and use a proportional shrinking-expanding proposal mechanism to update its value. All other details remain the same and are described elsewhere (Mitchell and Rabosky 2017).

### Simulation study

We conducted a simulation study to assess how well the method estimates branch specific rates of evolution and infers the presence of rate-shift events. We also evaluated whether the estimated rates of trait evolution accurately reconstruct ancestral character states.

#### (1) Choice of phylogeny

To carry out the simulations, we generated 100 phylogenies having between 50 and 1,000 tips by randomly sampling internal nodes from the 3,962-tip ultrametric squamate reptile phylogeny of Pyron and Burbrink (2014). Nodes were assigned weights such that all sizes (measured as the number of living descendants) of extracted clades had an equal probability of being selected. We chose to select subsets of a large empirical phylogeny, rather than simulated phylogenies, to introduce more realistic distributions of branch lengths than might be obtained using simple tree simulation models (e.g. Yule or constant-rate birth-death models).

#### (2) Simulating trait evolution

For each phylogeny, we simulated evolution of a binary state trait 10 times using 2 different procedures. In the first case, we determined the number of rate-shift events to place on the tree by drawing a random integer from a Poisson distribution with a rate of 1. We determined the locations for these rate-shifts by selecting that number of internal nodes randomly without replacement, again using weights that gave all sizes of subtrees an equal probability of being chosen, and choosing a uniform random point along each chosen node’s branch. We chose the transition rate for each rate-regime by drawing a random number from a log-normal distribution with a mean of log0.01 and a standard deviation of 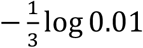. This corresponds to a log-normal distribution with a spread that position rates of 0.0001 and 1 three standard deviations below and above the mean, respectively, and these values were chosen simply for their potential to generate datasets having a range of phylogenetic signals. The second case was identical to the first except that we allowed transition rates to be asymmetric. The degree of asymmetry was determined by drawing a random number from a log-normal distribution with a mean of log 1 and a standard deviation of 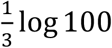. This corresponds to a log-normal distribution centered on unbiased transition rates with a spread that positions a 100-fold bias in transition rates 3 standard deviations from the mean.

#### (3) Information content of rate-shift events

Due to the stochastic nature of the simulations we expected rate-shift events to vary in their degree of detectability. To quantify this, we followed Rabosky et al. (2017) and calculated the “information content” of each rate-shift, which is a measure of how strongly the data support a model with rate variation over a model with no rate variation. For each simulated trait distribution, we optimized the value of the transition rate parameter to maximize the likelihood of the full data under a model with no rate-shifts. We denote this maximum-likelihood parameter estimate by *θ_CR_*. Next, we denote by *D_i_* the trait data contained in the subtree formed by the set of all nodes and branch segments belonging to the *i*-th rate-regime and by *θ_i_* the transition rate that generated those data. We calculated the information content of the *i*-th rate regime as the difference in log likelihoods of *D_i_* under the two parameter sets,

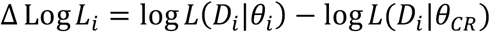

Where *L*(*D_i_*|*θ_i_*) denotes the likelihood of the data in the *i*-th rate regime given the generating parameter, and *θ_CR_* indicates the transition rate obtained by maximizing the likelihood for the full data under a model with no rate-shifts. If the data support a model with rate-shift events, the ΔLog *L* statistic must be greater than 0. In general, we expect that this statistic must be substantially greater than 0 for BAMM to detect a rate-shift event. If ΔLog *L* is expressed using the Akaike Information Criterion it can be rewritten as ΔLog *L* = ^S^/_2_ + *k*, where *S* is the difference in AIC scores needed to accept a model with an additional rate-shift and *k* is the number of extra parameters required to fit an additional rate-shift (in our implementation *k* = 2, corresponding to the location of the rate-shift and its rate parameter). If *S* ≥ 0 is interpreted as support for a model with an extra rate-shift, the minimum ΔLog *L* needed to detect an event is 2.

### BAMM analysis

We analyzed each simulation with BAMM using a single Markov chain that ran for ten million generations with Λ set equal to 1. The transition rates governing trait evolution need starting values before BAMM can proceed. For each simulation, we divided the total number of observed character state transitions by the summed branch length of the simulation’s phylogeny and used this value for the initial transition rate and for the median of the exponential prior placed on the transition rate. This number is unavailable in empirical datasets but will be close to the number implied by parsimony when rates of character evolution are low.

### Performance assessment

For each dataset simulated under a symmetric or asymmetric rate-shift model we assessed BAMM’s ability to estimate rates of trait evolution and to detect rate-shift events. We performed each assessment using the estimated posterior distribution after discarding the first ten-percent of samples.

#### (1) Estimating the rate of trait evolution

To determine how accurately BAMM infers rates of trait evolution we scaled each branch length to correspond to its average estimated rate of trait evolution. We performed this branch-by-branch calculation for each sample in the simulated posterior distribution using the BAMM-estimated transition rate parameters and assigned each branch an overall length equal to,

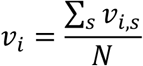

Where *v_i,s_* denotes the rate of evolution of branch *i* in the *s*-th posterior sample, and *N* indicates the number of samples in the posterior distribution. Using these values, we computed the tree-wide proportional branch rate error for a simulation as,

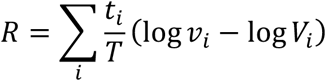

Where *V_i_* denotes the true rate of evolution of branch *i*, and the summation is taken over all branches in the phylogeny. The error of the *i*-th branch estimate was weighted by the proportional contribution of its branch length, *t_i_*, to the sum of all branch lengths, *T*, in the phylogeny. For unbiased transition rate estimates this equation is equal to 0.

#### (2) Detection of rate-shift events

For each simulated rate-shift event, we computed the mean detection accuracy as 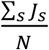, where *J_s_* indicates the detection accuracy in the *s*-th posterior sample and *N* indicates the number of samples in the posterior distribution. Letting *C_s_* denote the set of rate-shift events detected by BAMM in the *s*-th posterior sample, we measured the detection accuracy of a generating rate-shift event as,

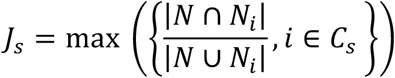

Where *N* is the set of tips descended from the generating rate-shift event and *N_i_* is the set of tips descended from the *i*-th rate-shift detected by BAMM in the *s*-th posterior sample. A value of 1 occurs when BAMM identifies the precise node that corresponds to the generating rate-shift event, and a value of 0 occurs when BAMM fails to identify the generating event at all. Intermediate values occur when BAMM identifies rate-shift events that correspond to nodes above or below the node where the generating rate-shift event occurred.

#### (3) Empirical application

Finally, we analyzed an empirical dataset of mimetic coloration in snakes previously analyzed with several related methods (Davis Rabosky et al. 2016). Red-black banded coloration arises repeatedly among harmless colubrid snakes that occur in broad sympatry with dangerously venomous red-black banded coral snakes in the Neotropics and parts of the North temperate zone. The incidence of red-black banded coloration is particularly high among dipsadine snakes, a highly diverse clade of colubrid snakes that occur in local and regional sympatry with coral snakes across the Neotropics. Using the random local clock model (Drummond and Suchard 2010) and a Medusa-like (Alfaro et al. 2009) model of discrete trait evolution, Davis Rabosky et al. (2016) inferred elevated rates of mimetic color evolution within dipsadine snakes resulting from repeated independent origins of red-black banding coincident with the diversification of coral snakes in the Neotropics. We repeated their macroevolutionary rate analysis of this dataset using the method developed in this paper. We divided the number of parsimony-implied character state changes by the total branch length of the phylogeny to obtain a median rate for the transition rate prior and ran BAMM for 10 million generations with Λ set equal to 10. We repeated this analysis 3 times under 3 different transition rate prior specifications corresponding to a 2-, 5-, and 10-fold speedup over the median rate implied by parsimony.

## Results

### (1) Estimating the rate of trait evolution

Tree-wide proportional branch rate errors were low regardless of whether the data were simulated under a symmetric (mean = 0.09, median = 0.08) or asymmetric (mean = −0.27, median = −0.17) rate-shift model (Fig. 2). The correlations between estimated and true branch rates were high when data were simulated using a symmetric model (Pearson’s ρ = 0.64, Spearman’s ρ = 0.90) but were substantially noisier when data were simulated using an asymmetric model (Pearson’s ρ = 0.26, Spearman’s ρ = 0.60). When branch rates were multiplied by the temporal duration of each branch to convert them into the expected number of character state changes these correlations improved, particularly for the asymmetric model (*symmetric*: Pearson’s ρ = 0.66, Spearman’s ρ = 0.93; *asymmetric*: Pearson’s ρ = 0.37, Spearman’s ρ = 0.75). Spearman rank correlations were substantially higher than Pearson product-moment correlations for both models, indicating that relative branch rates are generally better estimated than absolute branch rates.

**Figure 2.**
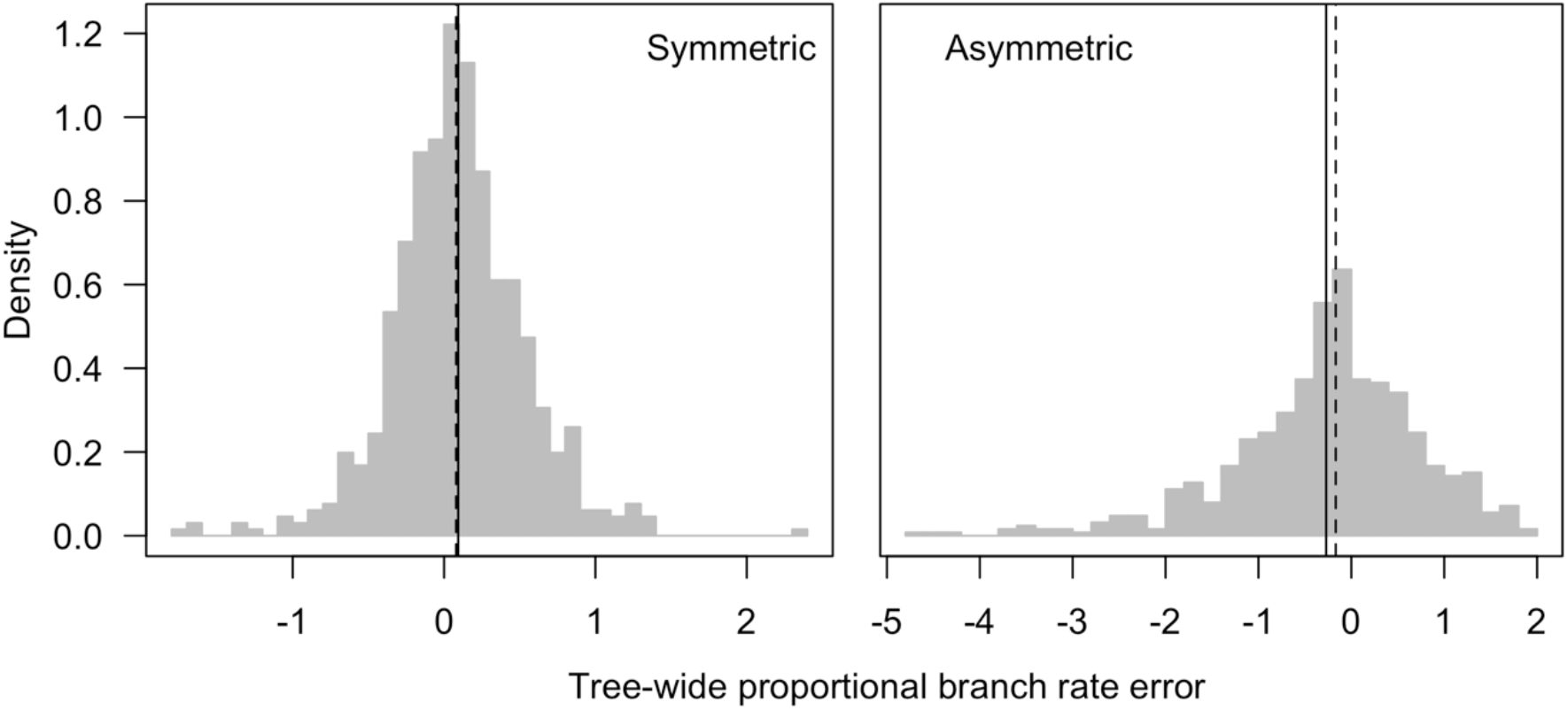
Tree-wide proportional branch rate error in BAMM-estimated branch rates when data are simulated under a symmetric rate-shift model (*left*) or an asymmetric rate-shift model (*right*). Branch rates are measured as the expected number of character transitions per million years. Tree-wide proportional branch rate error is a weighted sum of the logarithmic difference between estimated branch rates and true branch rates over all branches in a phylogeny. Errors on short branches are down-weighted relative to errors on long branches. Unbiased estimates have an error of 0, negative and positive values correspond to under- and over-estimation errors, respectively. Each histogram depicts the distribution of tree-wide proportional branch rate error over simulations with at least one rate-shift event. The vertical lines show the mean (solid) and median (dashed) of each distribution.

**Figure 3.**
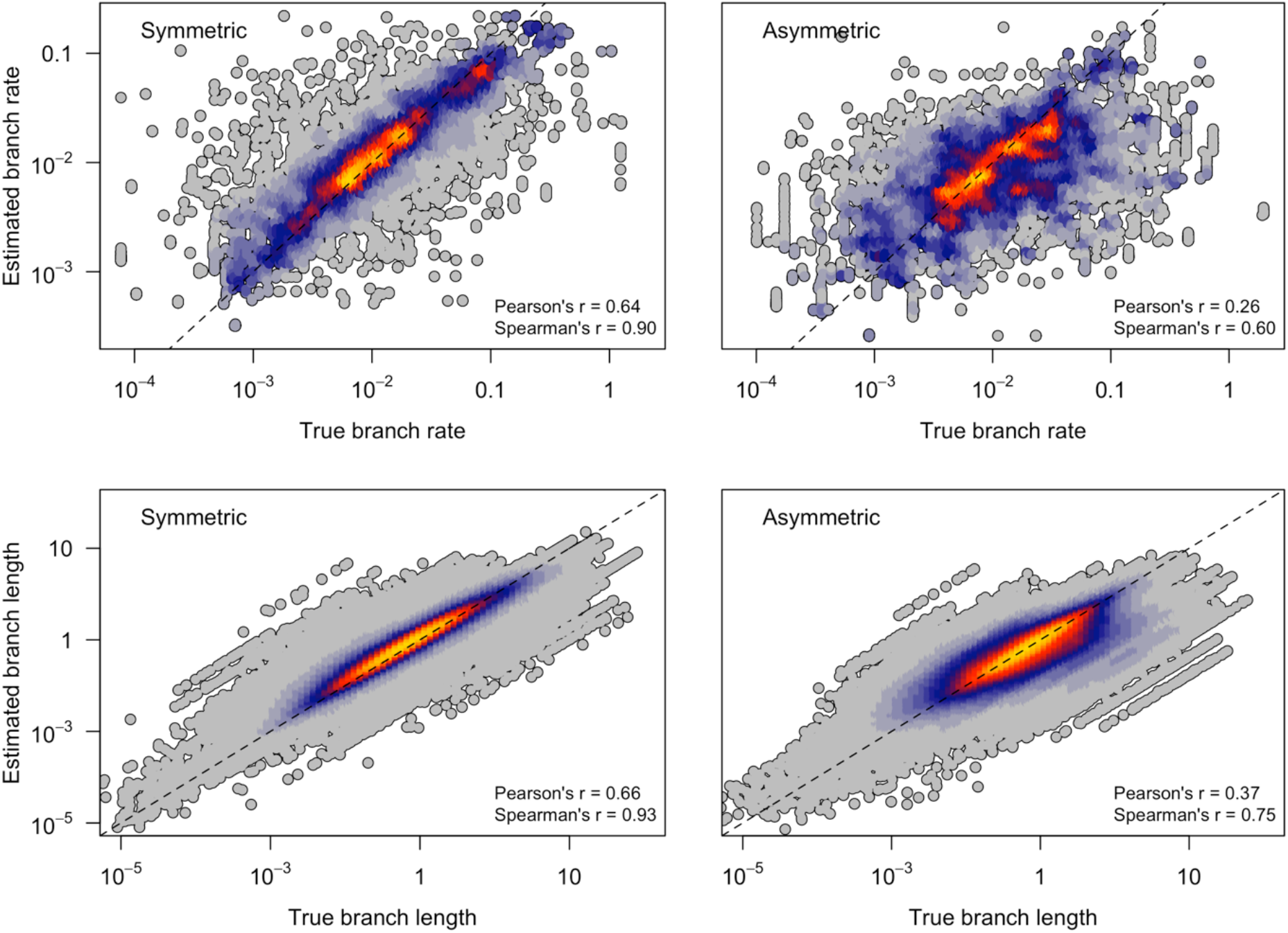
Correlation between true branch rates/lengths and BAMM-estimated branch rates/lengths when data are simulated under a symmetric rate-shift model (*left panels*) or an asymmetric rate-shift model (*right panels*). Branch rates (*top panels*) are measured as the expected number of character transitions per million years. Branch lengths (*bottom panels*) are measured as the expected number of character transitions occurring along the length of a branch. Each point depicts the average posterior rate estimate for a single branch of a phylogeny in one simulation, and the color of each point corresponds to the density of neighboring points (warm colors indicate high densities). Only branches from phylogenies with at least one simulated rate-shift are represented. The one-to-one line is dashed.

### (2) Detection of rate-shift events

BAMM’s ability to detect rate-shift events was generally low due to the limited information content of rate-shift events in the simulated data (Fig. 4). Despite relative rate differences between ancestral and derived rate-shift events that varied over 6 orders of magnitude, only 400 of 2045 simulated rate-shift events had an information content above 2, the theoretical minimum above which BAMM is expected to have power to detect them (see Discussion). This is due in part to the small average number of tips in simulated rate-regimes but also to the legitimate difficulty of simulating binary character data that reveal strong evidence for rate heterogeneity (cf. Fig. 4 rightmost panel). When BAMM has information to detect rate-shift events, however, it does reasonably well, and its performance improves monotonically as the information content of rate-shifts increases (cf. Fig. 4 leftmost panel). For example, BAMM detected the locations of 36 rate-shifts with an information content of at least 10 with a mean accuracy of 88%.

**Figure 4.**
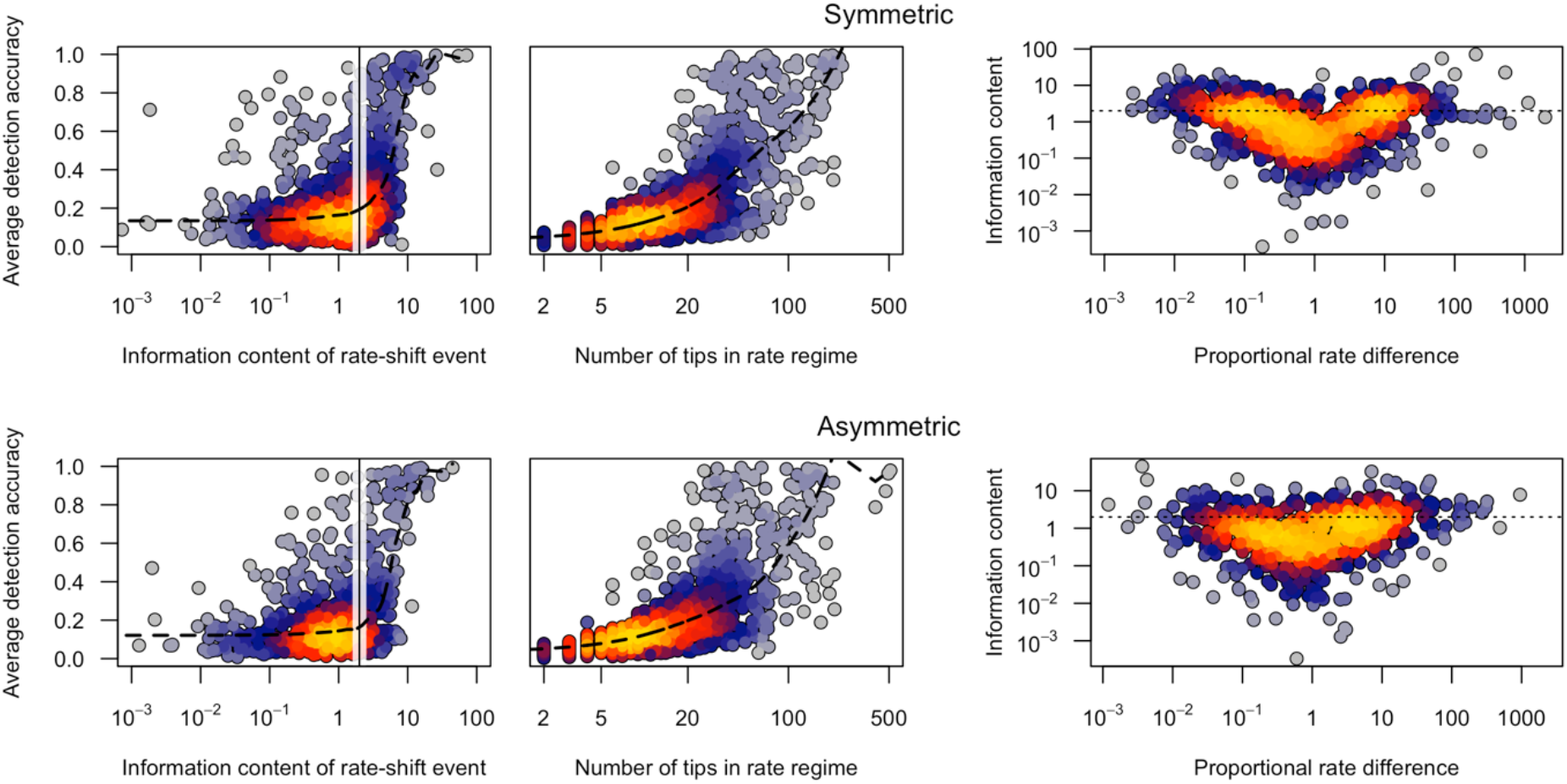
Rate-shift detection accuracy with BAMM when data are simulated under a symmetric rate-shift model (*top panels*) or an asymmetric rate-shift model (*bottom panels*). The average detection accuracy measures how accurately BAMM inferred a rate-shift’s location. The information content of a rate-shift event is a log-likelihood ratio that measures the likelihood of a given rate-shift event under the true parameters relative to the corresponding likelihood under a simple model where the rate is set to the whole-tree average. In the leftmost panels, the vertical line is the theoretical minimum log-likelihood ratio above which BAMM is expected to have power to detect a rate shift event, and it closely coincides with the upward inflection of the LOWESS regression lines (dashed). The rightmost panels plot an event’s information content against its proportional rate difference. The proportional rate difference of an event is the ratio of its rate of evolution to the ancestral rate preceding it. In all panels, each point represents a simulated rate-shift event, and the color of each point corresponds to the density of neighboring points (warm colors indicate high densities).

### (3) Empirical application

Analysis of the empirical dataset of red-black banded coloration in snakes largely recapitulated previous results showing an increased rate of trait evolution in Neotropical dipsadine snakes (but also revealed high rates in a clade of North temperate colubrine snakes) (Fig. 5). When analyzed under a strong, well-informed transition rate prior, rates of trait evolution ranged from a low of 0.00065 My^−1^ in basal snake lineages to a high of 0.0093 My^−1^ in dipsadine snakes, with an overall mean of 0.0024 My^−1^ that closely matched the overall rate of 0.002 My^−1^ obtained by dividing the number of parsimony-inferred state changes by the total branch length (cf. Fig. 5 leftmost panel). More liberal priors did not change this general picture, however rates of trait evolution in the upper tail of the estimated branch rate distribution tended to creep upward as the transition rate prior flattened. Flatter transition rate priors were associated with a higher number of posterior rate-shifts caused by the partitioning of single rate-shift events inferred under steeper transition rate priors into multiple smaller rate-shift events (i.e. lineages left out of these new rate-shift events fell back into the ancestral root regime). These new rate-shift events tended to have estimated rates of evolution that were higher than estimates made using steeper transition rate priors.

**Figure 5.**
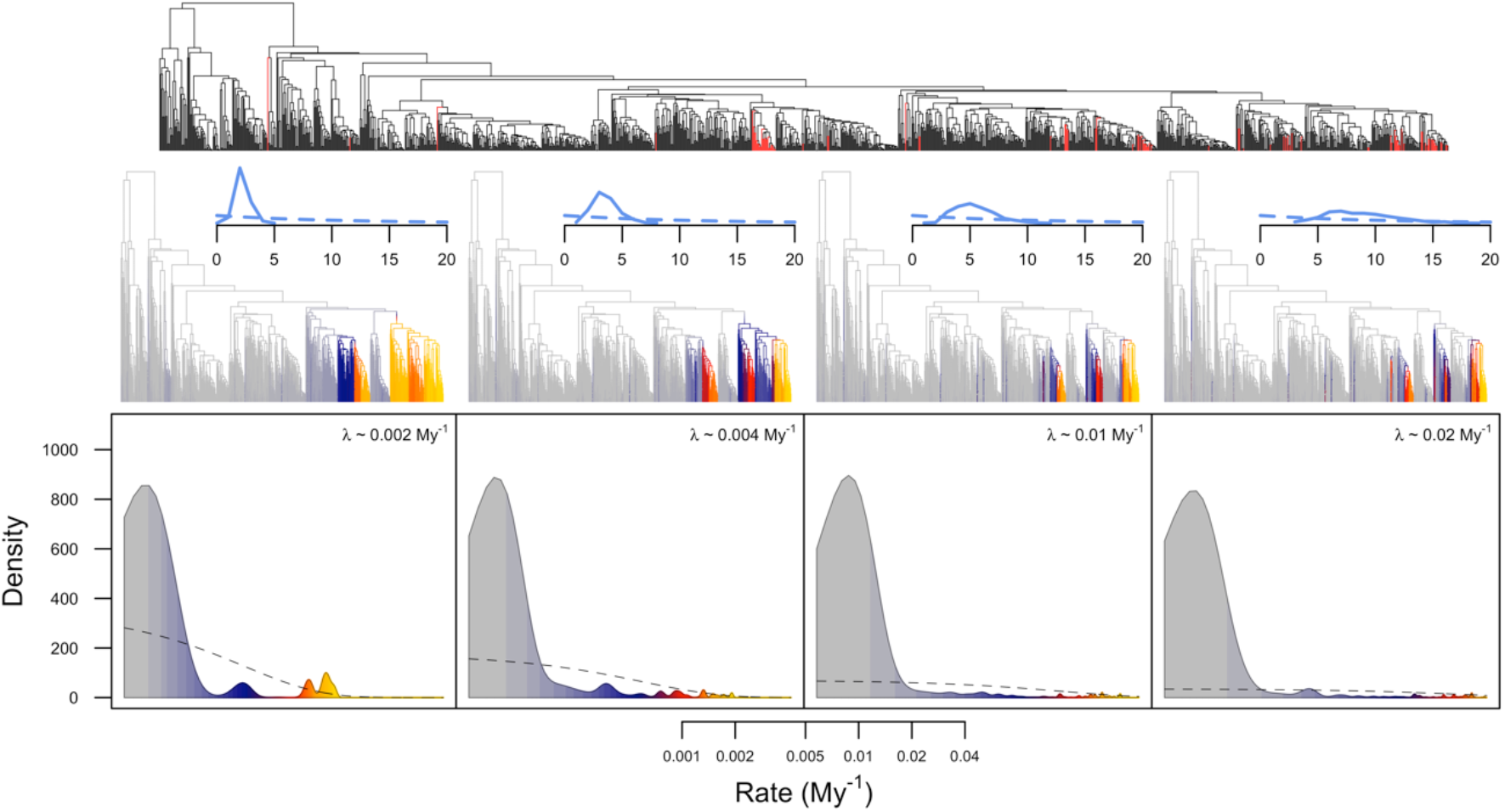
Evolution of mimetic coloration in snakes under four different transition rate priors in BAMM. A high rate of evolution of red-black banded mimetic coloration in snakes is inferred in Neotropical dipsadine snakes and North American colubrines, although precise locations of rate-shifts and rate estimates differ across prior specifications. In the lower panels, the *x*-axis is the average posterior rate of trait evolution (expected number of character state changes per million years) and the *y*-axis is a kernel density estimate of the branch-specific rate distribution over 4 different prior specifications. The dashed line shows the prior distribution of the rate of evolution and the prior median is indicated by the inset expression. Average posterior branch rates are mapped onto the phylogeny in the center panels. The line graphs above each phylogeny show the posterior distribution of the number of rate-shift events (solid line) in the credible shift set compared to the prior distribution (dashed line). Branches colored red in the topmost panel show the distribution of red-black banded mimetic coloration among tips in the empirical phylogeny. Phylogeny from Pyron and Burbrink (2014).

## Discussion

In this paper, we describe a Bayesian method, implemented in the BAMM software program, for studying among-lineage variation in the rate of evolution of a binary character. Overall, our results show that the method accurately infers rates of trait evolution and the presence and location of among-lineage evolutionary rate variation, even when simulated data violate model assumptions. Although the method performed well on many simulated data sets, we caution that overall power for inferring heterogeneous dynamics of single binary traits may be low.

The ability of the method to detect rate-shift events depends on the size of the clade belonging to a rate-regime but also on how much information the data contain with respect to the parameters of the rate-regime. In our simulations, we estimated this information content using a log-likelihood ratio that measures the likelihood of a given rate-shift event under the true parameters relative to the corresponding likelihood under a simple model where the rate is set to the whole-tree average. This information content can be surprisingly low even for large clades. In retrospect, this is not necessarily surprising because with only two character states there is a limit to how different data generated by two different rates can become. A 10- or even 100-fold rate increase will not be detectable if the ancestral rate is already high enough as to leave no phylogenetic signal. Similarly, a 10- or 100-fold rate decrease will not be detectable if the ancestral rate is already so low as to make character change highly improbable. In general, we expect detectability of rate-shifts to depend strongly on how distinct one rate’s phylogenetic signal is from another. This will have a strong stochastic component to it, and for binary data will likely have a low signal-to-noise ratio making detection of rate-shift events difficult.

While we did not explore prior sensitivity exhaustively in this study, the empirical results indicate that branch rate estimates (and ancestral state reconstructions by implication) are sensitive to the transition rate prior. In the empirical example, disagreements among the different priors occur in regions of phylogeny having elevated rates of evolution. This is not surprising given that the method works with only a single binary character and that the fast-evolving clades in the empirical phylogeny have relatively few taxa, but it does call for vigilance. A sensible rule of thumb is to treat with caution any result where the overall mean rate estimate disagrees strongly with the rate implied by parsimony. Encouragingly, the different prior specifications are in broad agreement on where relative rate differences occur in the empirical phylogeny despite disagreements over absolute rate estimates. The results also indicate that the transition rate prior interacts with the prior on the number of rate-shifts, suggesting that the method’s ability to estimate the precise location of a rate-shift and its ability to estimate the associated rate of evolution may trade-off. Finally, the combination of a prior with a high number of expected rate-shift events and a relatively flat transition rate prior can lead to an apparent abundance of single tips with derived character states having elevated rates of evolution. This is because dropping a high rate of evolution on such a branch entails no penalty. It makes the derived state more probable and with only a single lineage does not suffer from the likelihood penalty that a larger clade fixed for a derived state would suffer from if given an elevated rate. Users should remain alert to this scenario and treat its presence as an indication that the priors are exerting an undue influence. For all these reasons, we recommend the use of a strong, well-informed prior on the transition rate and setting the median rate of the prior equal to the parsimony-implied rate seems like a sensible choice.

A fundamental difficulty facing macroevolutionary models of discrete character evolution is that the data contain low information content with respect to rates of evolution since actual events of character state change are not directly observed. This challenge is exacerbated further in asymmetric Markov models which must estimate the rate of evolution while simultaneously inferring the equilibrium frequencies of the character states. From this fact alone we should expect asymmetric Markov models to have lower information content with respect to transition rates than symmetric Markov models, which a priori assume equilibrium frequencies of character states are equal. In a discrete-time version of the Markov model used by this and many other studies, Sanderson (1993) has shown that maximum likelihood parameter estimates are intrinsically biased upwards for an asymmetric model with two character states but are unbiased for a symmetric model. The extent to which these conclusions apply in continuous-time or generalize to more than two states is not known to us but warrants further study.

One consequence of the low information content of binary data are ancestral state reconstructions that may appear nonsensical when performed with asymmetric models. For example, Pagel (1999) presented a comb phylogeny in which every tip in the clade was fixed for one of two possible character states and showed that an asymmetric Markov model reconstructed the root as belonging to either state with equal probability. From an asymmetric model’s perspective, a clade that is almost entirely fixed for a single character state is likely to have been generated by a process with a very high transition rate toward the majority state and a very low transition rate away from it, which makes the tip states mostly independent of the ancestral states because the asymmetry in rates will yield the same outcome at the tips regardless of the assignment of states to internal nodes. By contrast, from a symmetric model’s perspective the clade simply has a very low rate of evolution and the tip states resemble their ancestors as a result. In fact, transition rates are nearly always biased in the direction of the state appearing most frequently among the tips of the tree (Nosil and Mooers 2005; Maddison 2006). This means that in an analysis that infers a high rate of transition from state 0 to state 1 and low rate of transition from state 1 to state 0, it will sometimes be the case that a majority of the evolutionary transitions were from state 1 to state 0. This runs counter to intuition and indicates the need for caution when using transition rates to infer directionality in the history of trait evolution (Nosil and Mooers 2005; Goldberg and Igic 2008).

## APPENDIX

*Derivation of likelihood equations (1) and (2)*

Begin with the differential equations,

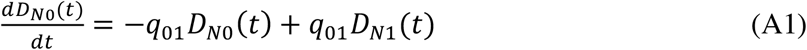

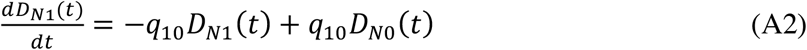

By solving (A1) for *D*_*N*1_(*t*) and equating its derivative to (A2) we can form the second-order differential equation,

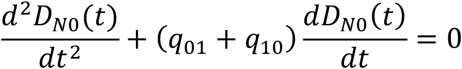

Which, after finding the roots of its auxiliary equation, has the general solution,

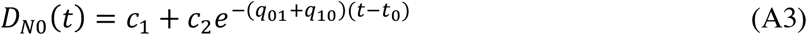

Where *t*_0_ is a starting time on node *N*’s branch, *c*_1_ and *c*_2_ are constants, and *t* is larger than *t*_0_ but smaller than the time at the base of node *N*’s branch.

To solve for *c*_1_ note that at our initial condition when *t* = *t*_0_,

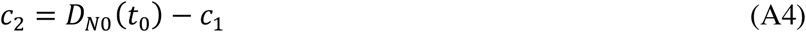

Furthermore, differentiating equation (A3) gives,

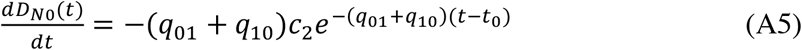

And equating (A1) with (A5) and setting *t* = *t*_0_ yields, after some algebra,

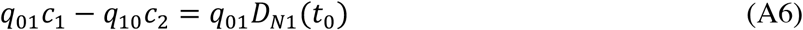

By substituting the right-hand side of (A4) for *c*_2_ in (A6) and solving for *c*_1_ we find,

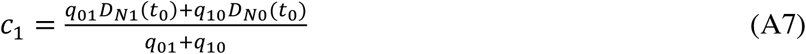

Finally, by substituting the right-hand side of (A7) for *c*_1_ in (A6) and solving for *c*_2_ we find,

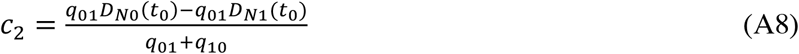

Thus, equation (A3) is equivalent to equation (1) after substitution of (A7) and (A8), noting that 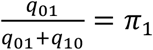 and that 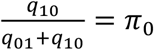. An equivalent derivation of equation (2) is performed by starting with *D*_*N*1_(*t*) = *c*_1_ + *c*_2_*e*^−(*q*_01_+*q*_10_)(*t*–*t*_0_)^. When 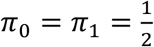 as in the BAMM implementation this is simply a special case of the JC69 model (Jukes and Cantor 1969) with two states.

